# Spatial expression evolution of duplicated genes post-whole genome duplication in cultivated tomato

**DOI:** 10.64898/2026.10.28.762560

**Authors:** Thilanka Ranaweera, Rachel E. Kerwin, Huan Chen, Robert L. Last, Peipei Wang, Shin-Han Shiu, Melissa D. Lehti-Shiu

**Affiliations:** Department of Plant Biology, Michigan State University, East Lansing, MI, 48824, USA; DOE Great Lakes Bioenergy Research Center, Michigan State University, East Lansing, MI, 48824, USA; Department of Biochemistry and Molecular Biology, Michigan State University, East Lansing, MI 48824, USA; 4Kunpeng Institute of Modern Agriculture at Foshan, Agricultural Genomics Institute at Shenzhen, Chinese Academy of Agricultural Sciences, Shenzhen, Guangdong 518124, China; Department of Computational Mathematics, Science, and Engineering, Michigan State University, East Lansing, MI, 48824, USA

**Author notes:** The author responsible for distribution of materials integral to the findings presented in this article in accordance with the policy described in the Instructions for Authors (https://academic.oup.com/plphys/pages/General-Instructions) is Shin-Han Shiu.

**Keywords:** Ancestral State Reconstruction, Expression Divergence, Organ-Specific Expression, Post-WGT Paralogs, Solanaceae-Specific Whole Genome Triplication, Spatial Dependence of Expression, Syntenic Gene Blocks

## Abstract

The mechanisms by which whole genome duplication contributes to expression divergence of duplicate genes, which can impact the evolution of form and function, has been a long-standing question in evolutionary biology. We compared expression patterns of paralogs derived from the most recent Solanaceae-specific genome triplication event across 23 tissues and inferred ancestral expression states to investigate patterns of expression evolution. While the partitioning of ancestral expression states is traditionally considered the second most common (after the gene loss) trajectory of duplicate expression evolution, we found that the ancestral expression states tended to be retained by most (∼66%) duplicates, with ∼6% of gene lineages experiencing expression gain events and ∼20% experiencing expression losses. The genes that gained expression in fruit tissues are significantly enriched with functions associated with fruit quality. Notably, asymmetrical expression divergence is prevalent among duplicates, especially when ancestral genes were either not expressed or were highly expressed. Our study demonstrates the utility of integrating paralog expression comparisons with phylogenetic analyses to document the potential evolutionary history of spatial expression patterns and gain insight into how expression divergence contributes to the evolution of function across tissues.

## INTRODUCTION

A key objective in evolutionary biology is to develop a mechanistic understanding of how gene duplication contributes to the emergence of novel adaptive traits. In particular, whole-genome duplication (WGD), which doubles all genetic material, provides opportunities for gene functional diversification, specialization, and novelty (Ohno, 1970; Ren *et al*., 2018). While the fate of most duplicates is eventual loss (Lynch and Conery, 2000; Maere *et al*., 2005), many are retained and can be co-opted to create new pathways or subnetworks that contribute to evolutionary innovation (True and Carroll, 2002).

Over the past two decades there has been significant progress in understanding the evolutionary trajectories of WGD duplicates, with several models proposed to explain duplicate gene retention (Liu and Adams, 2007; Hovav *et al*., 2008; Liu, Baute and Adams, 2011a; Arsovski *et al*., 2015; Coate *et al*., 2020; Garber *et al*., 2024). The neofunctionalization model (Ohno 1970), in which one duplicate retains the ancestral gene function while the other acquires a novel function, has been demonstrated to play an important role in development of novel structures (He and Saedler, 2005; de Martino *et al*., 2006; Zhang *et al*., 2015), metabolic innovations (Durbin, McCaig and Clegg, 2000; Fiesel *et al*., 2022; Kerwin *et al*., 2024), and stress tolerance (Zou *et al*., 2009; Yang *et al*., 2014; Lee and Adams, 2020). Another widely accepted model is the duplication-degeneration-complementation (DDC) model (Force et al., 1999; Lynch and Force, 2000), in which ancestral functions are partitioned among paralogs, a process known as subfunctionalization. Consistent with this model, studies have shown that a large proportion of retained duplicates show evidence of partitioning of ancestral expression states between copies. For example, in *Arabidopsis thaliana* about 45% of paralogous pairs show partitioning of ancestral expression states across organs, and higher levels of partitioning are observed among WGD duplicated transcription factors (Duarte *et al*., 2006; Panchy *et al*., 2019). Several other models, such as the Gene Balance Hypothesis (Birchler *et al*., 2001; Veitia, 2005), the Increased Gene Dosage Hypothesis (Seoighe and Wolfe, 1999; Kondrashov and Kondrashov, 2006), and the Functional Buffering Model (Chapman *et al*., 2006), have been proposed to explain gene retention following duplication events.

Gene expression is a major determinant of gene function (Romero, Ruvinsky and Gilad, 2012); therefore, studying how expression evolves after WGD can provide insight into the functional divergence of duplicated genes (Adams *et al*., 2003; Liu and Adams, 2007; Ha, Kim and Chen, 2009; Zou *et al*., 2009; Panchy *et al*., 2019). Moreover, investigating the evolutionary processes underlying expression divergence of WGD duplicates at a genome-wide scale can provide insights into the genetic and developmental mechanisms underlying trait evolution (Duarte *et al*., 2006; Liu and Adams, 2007; Ha, Kim and Chen, 2009; Yang *et al*., 2014). Currently, two main approaches are commonly employed to study expression evolution at a genome-wide scale: (1) pairwise studies, which examine expression divergence between paralogs within a species (Gu *et al*., 2002; Krylov *et al*., 2003; Makova and Li, 2003), and (2) phylogenetic comparative methods, which compare expression across orthologous genes to infer expression dynamics relative to ancestral states (Oakley *et al*., 2005; Brawand *et al*., 2011; Rohlfs and Nielsen, 2015). However, phylogenetic comparative methods, which are commonly used in combination with ancestral state reconstruction, face challenges. These include ortholog assignment uncertainty due to lineage-specific duplications or losses and unreliability of ancestral state estimations from deep phylogenies (Martins, 1999; Koonin, 2005; Oakley *et al*., 2005; Hahn, 2007). Pairwise comparisons, in contrast, directly measure the difference between paralogs and avoid technical biases arising from cross-species comparisons. Nevertheless, without phylogenetic context, pairwise methods cannot reliably infer how functional divergence has occurred because the ancestral states are unknown (Oakley *et al*., 2005; Conant and Wolfe, 2008). To overcome these limitations, we applied an integrated approach that combines pairwise comparison methods inferring expression divergence from extant paralog expression states, and phylogenetic comparative methods modeling the ancestral expression states, to investigate how gene expression evolves after whole genome duplication in cultivated tomato (*Solanum lycopersicum*) (Sato *et al*., 2012).

We chose tomato as our model system for two reasons. First, tomato is a model for studying fleshy fruit development, and there is substantial understanding of the genetic mechanisms underlying fruit size, shape, color, and flavor traits (Lin *et al*., 2008; Sato *et al*., 2012; Kim *et al*., 2018; Quinet *et al*., 2019). Second, the Solanaceae lineage has undergone two whole genome triplication (WGT) events since it diverged from the most recent common ancestor of angiosperms: one shared with the Rosids and Asterids, and the other exclusive to the Solanaceae lineage (hereafter referred to as Sol-WGT) (Sato *et al*., 2012; Huang *et al*., 2022). These WGTs, followed by natural selection shaping the retention and diversification of duplicate genes, and later directional selection through breeding, have driven the expansion of gene families involved in cell wall metabolism, hormone signaling, and specialized metabolism, ultimately contributing to the evolution of traits such as fruit texture, flavor, and nutritional composition (Flagel and Wendel, 2009; Sato *et al*., 2012; Rajewski *et al*., 2022). Furthermore, the retention of fruit-specific transcriptional regulators post WGT has been linked to the emergence of developmental and ripening processes, demonstrating how selection of WGT paralogs during domestication has shaped the tomato genome (Sato *et al*., 2012). Therefore, by characterizing the expression evolution of these recent duplicates, we can identify the specific modes of functional divergence that have catalyzed the emergence of novel traits within the Solanaceae lineage.

In this study, we investigated how expression patterns diverged for paralogous genes derived from the Sol-WGT in cultivated tomato using two approaches: (1) comparing extant expression profiles between Sol-WGT duplicates across tomato tissues, and (2) comparing extant gene expression states with ancestral ones (Fig. 1A). Here we demonstrate different expression divergence patterns in multiple tissues of tomato by comparing the ancestral and extant expression states. We show that expression is likely to evolve in a coordinated manner among tissues, especially those from similar organs. Our results provide insights into the expression divergence of duplicates after whole genome duplication and how it shapes adaptive trait evolution.

**Figure 1:**
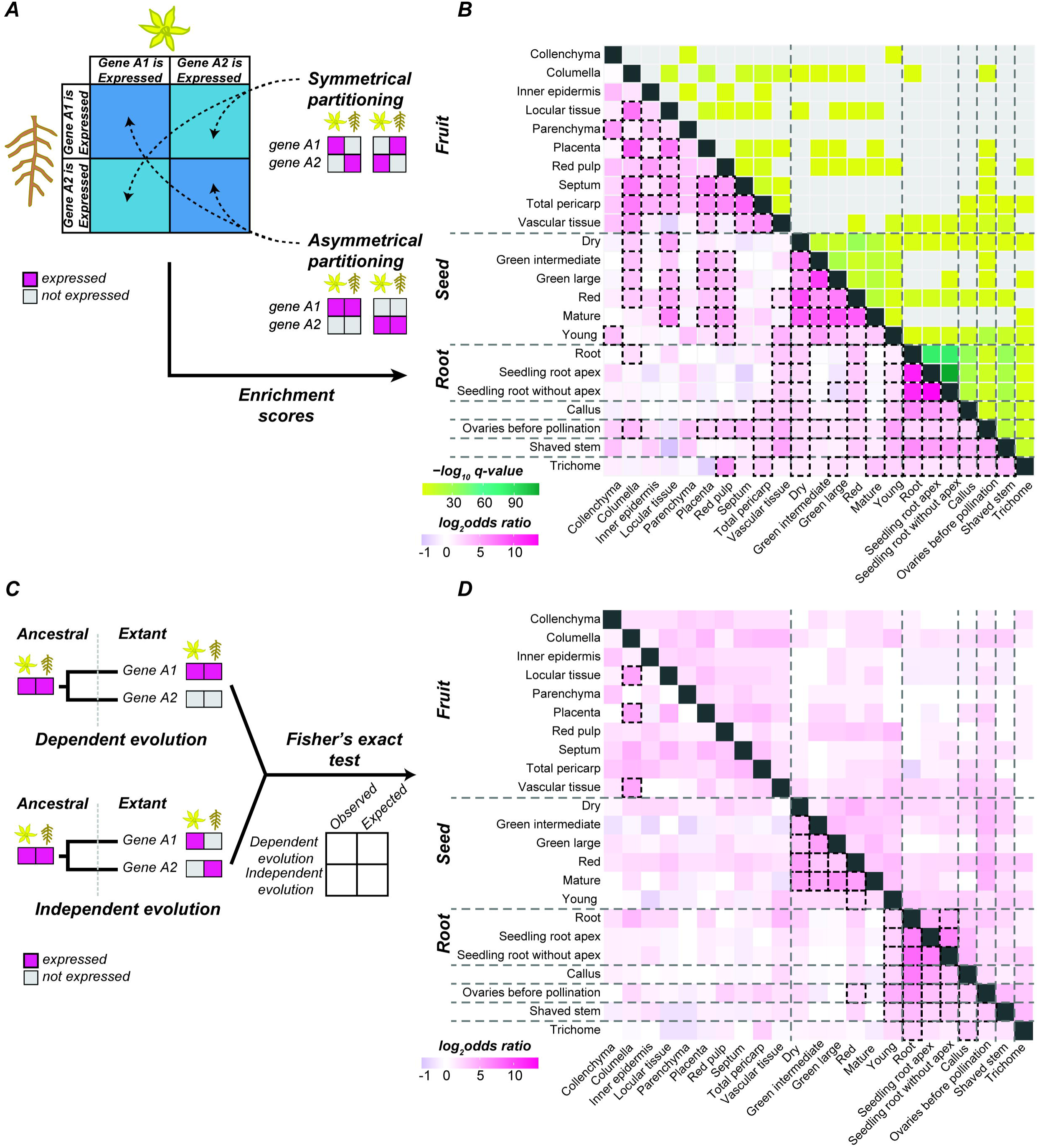
Identification of genes derived from the Solanaceae-specific whole genome triplication (WGT). **(A)** Schematic diagram illustrating the distinctions between “Expression divergence considering both ancestral and extant expression” and “Expression divergence considering only extant gene expression” as used in this study. The phylogenetic tree depicts the evolutionary history of Gene A, which gave rise to Gene A1 and Gene A2 following a whole genome duplication (WGD) event. Tissues in which extant or ancestral genes are expressed are in magenta, while tissues with no expression of a given gene are colored gray. **(B)** Ks (synonymous substitution rate) plots for delineating major WGT events. The gray line represents the median Ks distribution of all syntenic blocks in the tomato genome. Mixed modeling reveals three distinct distributions: two normal distributions, one (shaded blue) corresponding to the Sol-WGT event and one (orange dashed line) associated with the γ WGT event, and an exponentially decaying distribution (violet dashed line) reflecting more recent episodic duplication events. Two vertical dotted lines indicate the Ks thresholds (t1 and t2) used to determine syntenic blocks derived from the Sol-WGT event. **Alt Text:** Diagrams of a phylogenetic tree and tissue expression alongside three synonymous substitution rate plots that use mixed modeling and color-coded distributions to identify whole genome triplication events.

## MATERIALS AND METHODS

### Collection and processing of transcriptomic data and calculation of absolute expression levels and differential expression

Total transcriptomic data for 46 different tissue types of *S. lycopersicum* (Table S1) were downloaded from the National Center for Biotechnology Information (NCBI) Sequence Read Archive (SRA). Tomato (Cultivar: Heinz1706) reference genome (build 4.00) gene annotations (ITAG 4.0 release) and proteome data (ITAG 4.0 release) were downloaded from Solanaceae Genomics Network (https://solgenomics.net/). The raw RNA sequencing (RNA-seq) reads were initially converted to fastqc format using fastq-dump in SRA-Toolkit (https://github.com/ncbi/sra-tools). The read lengths and the quality of the raw sequences were determined using FastQC (Andrews, 2010). To get the clean reads, the raw RNA-seq reads were trimmed using Trimmomatic V. 3.39 (Bolger et al., 2014); all settings were the default except for LEADING:20, TRAILING:20, SLIDINGWINDOW:4:20, and MINLEN:36. The paired-end and single-end sequences were trimmed separately, and the adapters were clipped using Illumina paired-end and single-end adapter sequences, respectively. The clean reads were mapped to the tomato genome using Tophat2 (Kim *et al*., 2013), and read counts were obtained using HTseq (Anders, Pyl and Huber, 2015). The transcript abundance was measured in transcripts per million (TPM) by processing the count data using the Scatter package (Version 1.34) implemented in R (McCarthy *et al*., 2017). All the RNA-seq data were processed using a custom RNA-seq data processing pipeline (https://github.com/ShiuLab/RNA-seq_data_processing.git).

### Analysis of expression divergence between post-WGT duplicates

For all the post-WGT duplicates (see supplementary methods for the identification of paralogous genes derived from the Solanaceae-specific WGT event), the number of tissues in which each gene is expressed (TPM ≥ 2) was calculated (see supplementary methods for determination of the cutoff for gene expression; Fig. S1A). To determine how the ancestral expression pattern was partitioned among paralogs, the number of tissues in which each copy of a paralogous gene pair was expressed was calculated. Then, for all paralogous gene pairs, the frequency of different tissue expression combinations was determined (e.g., if one paralog was expressed in 10 tissues and the other in 5, the occurrence of the [10, 5] combination among all paralogous pairs was calculated). A frequency table was then generated to capture all possible combinations of tissue expression patterns between genes for all the syntenic paralogous gene pairs (Fig. 2A and B). Next, deviation of the observed expression divergence between paralogous gene pairs from that expected randomly was determined by conducting simulations to model expected scenarios (Fig. 1A and C). During each iteration of the simulation, the observed extant expression states for tissues were randomly swapped between the paralogs within each specific tissue type. For example, if one gene of a paralogous pair was expressed in trichomes while the other was not, during the random swap, both genes had an equal probability of being assigned the expression state, while maintaining the total number of tissues in which the pair was expressed. This simulation was run for 10O iterations. At the conclusion of the simulation, the frequency distribution (with 10O instances) of tissue expression combinations between paralogous gene pairs was recorded. These simulated distributions were then compared to the observed data to calculate a Z-score reflecting how much observed frequencies of different expression combinations differed from random expectations (custom code available at GitHub: https://github.com/ShiuLab/Manuscript_Code/tree/master/2025_Tomato_expression_evolution).

**Figure 2:**
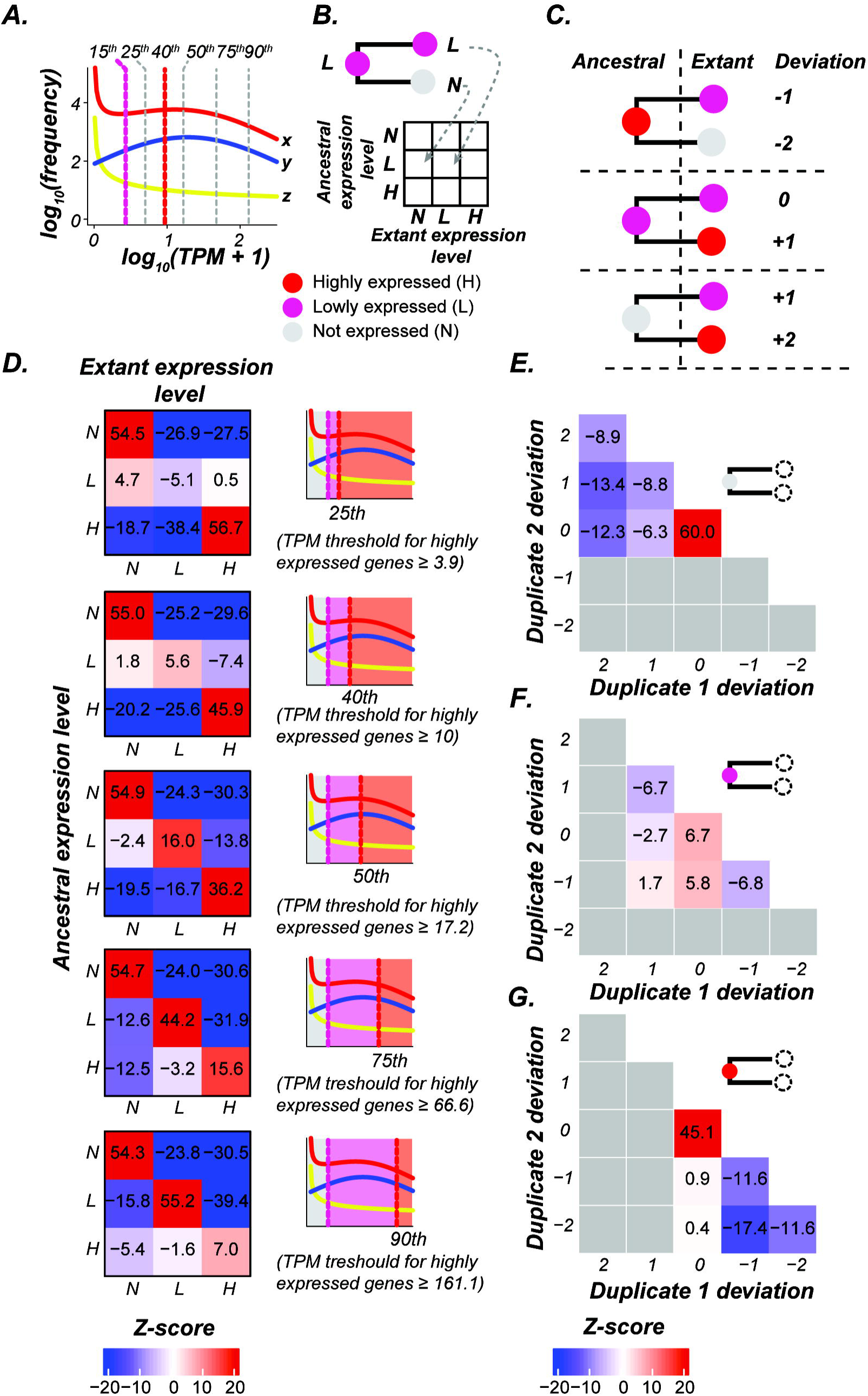
**Analysis of divergence in expression breadth using extant expression data**. **(A)** Illustration of how expression divergence between paralogous gene pairs is calculated as a Z-score. First, a frequency table is generated to capture all possible combinations of tissue expression patterns for paralogous gene pairs, based on how many tissues in which each gene is expressed (TPM ≥ 2). For example, if gene A1 is expressed in four tissues and gene A2 is not expressed in any, the corresponding cell in the frequency table specifying this pattern is incremented by one. The expected expression divergence scenarios are estimated as detailed in the “Analysis of expression divergence between post-WGT duplicates” in the Materials and Methods section. Left panel: observed frequency, right panel: Z-score of the observed frequency. **(B)** Observed frequencies for all combinations of paralogous pair expression in 23 tissues. Darker green indicates higher frequency. Genes in a pair are randomly assigned the labels gene 1 and 2. **(C)** Heatmap showing Z-scores calculated for all paralogous gene expression breadth scenarios. Color represents the sign of the Z-score, and intensity indicates the magnitude. Black boxes are the scenarios that did not occur in the simulation. **Alt Text:** Statistical schematics and color-coded heatmaps illustrating the frequency and Z-score calculations for expression breadth divergence between paralogous gene pairs across twenty-three different plant tissues.

### Determination of the spatial dependence of extant expression of post-WGT duplicates

To determine the association of expression patterns between paralogous gene pairs across different tissue types (spatial dependence), pairwise tissue comparisons were performed. For each tissue pair, paralogs exhibiting expression divergence (in the cases where both paralogous genes are not expressed in the same tissue) were classified as exhibiting symmetrical or asymmetrical expression divergence. For example, when comparing trichomes and root hairs, symmetrical divergence was defined as one paralog being expressed in trichomes but not in root hairs, while the other paralog showed the opposite pattern (expressed in root hairs but not in trichomes). Asymmetrical divergence was defined as one paralog being expressed in both tissues while the other was not expressed in either. We then used Fisher’s exact test (Fisher, 1922) to test the significance of deviations from the null hypothesis that expression divergence between paralogous gene pairs is symmetrical. P-values were calculated, with log odds ratios serving as enrichment scores, for all possible tissue pair combinations. This analysis was conducted on the entire set of paralogous genes, as well as subsets of paralogous gene pairs that were not expressed in more than 10, 5, or 2 tissues (custom code available at GitHub: https://github.com/ShiuLab/Manuscript_Code/tree/master/2025_Tomato_expression_evolution).

### Analysis of expression divergence of the Sol-WGT duplicates after the WGT event

To test how ancestral gene expression diverged among post-WGT paralogs (see supplementary methods: “Identification of paralogous genes derived from the Solanaceae-specific WGT event” for identification of paralogous genes), the reconstructed ancestral expression states were compared with the extant expression states for each tissue type (see supplementary methods sections “Phylogenetic tree reconstruction” and “Ancestral expression state reconstruction”; Fig. S2A-F). Ancestral expression levels at the Solanaceae-specific WGT event were retrieved from the phylogenies generated for each syntenic paralog–tandem duplicate group. Both the extant and ancestral expression abundances were classified into three categories: not expressed, expressed, and highly expressed. As previously described, a TPM threshold of two was used to distinguish between expressed and non-expressed genes (see supplementary methods for determining absolute expression threshold). Highly expressed genes were identified using different percentiles of a modeled normal distribution of absolute expression levels (Fig. S1A). A gene was considered expressed if its expression level (TPM) was ≥ 2 and less than the highly expressed threshold, while a gene was classified as highly expressed if its expression exceeded the highly expressed threshold. To assess the impact of varying thresholds on the interpretation of expression partitioning, we used five different highly expressed thresholds (TPM = 3.9, 10, 17.2, 66.9, 161.1).

Afterwards, the frequencies of different expression outcomes, gain, loss, or retention, of the observed scenarios were calculated. The observed scenarios represent the possible combinations of ancestral and extant expression states for each paralog in the syntenic paralog–tandem duplicate groups. For instance, a gene that was not expressed in the ancestral state but is expressed in the extant state would be categorized as a gene with gain in expression. To capture all possible combinations of ancestral and extant expression states, a frequency table was generated, detailing the occurrences of each of the nine possible transitions between ancestral and extant states (not expressed to not expressed, not expressed to expressed, not expressed to highly expressed, and so on). This allowed us to quantify the extent to which gene expression states had either been retained, gained, or lost for each paralog after the WGT event, across all tissue types examined.

A simulation was done to determine how the observed frequency of ancestral expression divergence deviated from the expected divergence. In the simulation, the extant expression states of syntenic paralogs were allowed to change independently from their ancestral expression states. In this model, regardless of the ancestral expression, the extant expression could take any expression state with equal probability. This simulation generated a frequency distribution for each scenario, which was then compared to observed frequencies. Z-scores were calculated to quantify the deviation of the observed expression divergence from the expected divergence.

Next, we compared the divergence of expression between syntenic paralogous gene pairs and their corresponding ancestor. This comparison assessed how the extant expression states (not expressed, expressed, or highly expressed) of syntenic paralogs deviated from the ancestral expression at the node corresponding to the WGT event. For this analysis, a threshold of TPM > 10 was used to classify genes as highly expressed, as it was the 40^th^ percentile of the true positive expression distribution, which eliminated 99% of the false positive expression (see supplementary methods: “Determination of the cutoff for gene expression”). The deviation was calculated based on changes in expression state from the ancestral to the extant state. A deviation score of +1 indicates a shift to a higher category, a deviation score of-1 indicates a shift to a lower expression category, and a score of 0 indicates no change. For example, in a particular tissue (e.g., root), if the ancestral gene was expressed, one of the paralogs was non-expressed, and the second paralog was highly expressed, the deviation would be recorded as-1 for the first paralog and +1 for the second paralog. A frequency table was constructed to capture all possible combinations of expression deviation between each paralog and the ancestral gene for all post-WGT paralogous gene pairs.

Next, to determine how the observed scenarios of expression deviation differed from expected scenarios, simulations were carried out in which the extant expression of each syntenic paralog was allowed to change independently from its ancestral expression, with each expression state (not expressed, expressed, highly expressed) assigned equal probability regardless of the ancestral state. From these simulations, we generated a distribution of expected frequencies for each scenario. The observed frequencies were then compared to the expected distribution to calculate Z-scores to quantify how significantly the observed deviations from the ancestral state differed from the random expectation. This analysis was done using custom codes (available at GitHub:https://github.com/ShiuLab/Manuscript_Code/tree/master/2025_Tomato_expression_evolution).

### Determining the association of expression between tissues considering ancestral gene expression

To evaluate the spatial dependence of the expression evolution of paralogous genes that arose from WGT events, we analyzed whether expression changes in syntenic paralog pairs were similar across tissue types. Syntenic paralog pairs in which one gene had gained or lost expression in a specific tissue (relative to the ancestral gene) while the other gene retained the ancestral expression pattern were selected for this analysis. To detect spatial dependence, we examined if the expression divergence was consistent between tissues for each paralogous pair. For example, if an ancestral gene was expressed in trichomes and one paralog retained this expression pattern while the other lost it, we checked if the same expression pattern occurred in another tissue, such as roots. If the paralogous gene retaining trichome expression also retained root expression, while the other paralog lost both, this was considered “dependent evolution” of expression between trichome and root tissues. Conversely, if one paralog lost root expression while the other retained it, this was classified as “independent evolution” of expression between tissues. A Fisher’s exact test was performed to determine if dependent or independent evolution of expression was more likely to occur (Fisher, 1922). The null hypothesis was that dependent and independent expression events between tissue pairs were equally distributed across paralogous gene pairs. This analysis included the entire set of paralogous gene pairs for which one gene had gained or lost expression in a specific tissue (relative to the ancestral gene) while the other gene retained the ancestral expression pattern. Including large numbers of paralogous gene pairs where one paralog is broadly expressed while other paralogous gene is narrowly expressed can inflate the “dependent evolution” scenarios. Thus, subsets of paralogous pairs that were expressed in no more than 10, 5, or 2 tissues were analyzed to test for potential bias from broadly expressed genes. Paralogous pairs for which the ancestral gene was not expressed (indicating a gain in expression) and those for which the ancestral gene was expressed (allowing analysis of both gain and loss events) were analyzed separately. Custom code for this analysis is available on GitHub: https://github.com/ShiuLab/Manuscript_Code/tree/master/2025_Tomato_expression_evolution.

## RESULTS AND DISCUSSION

### Identification of paralogs derived from the Solanaceae-specific WGT

To examine how gene expression evolved in the Solanaceae lineage, we compared the expression patterns of syntenic tomato genes retained from the Sol-WGT event (Fig. 1A). To do this, we first identified syntenic regions and estimated the age of duplicates within these regions by calculating the synonymous substitution rates (Ks) between syntenic paralogs (Ks_gene_) and the median Ks between syntenic gene blocks (Ks_block_).

In a Ks frequency distribution for all the pairs of genes/syntenic gene blocks, distinct peaks are indicative of paralogs arising from ancient large-scale duplication events (Schlueter *et al*., 2004; Cui *et al*., 2006). We observed that the frequency distribution drawn using Ks_block_ values (Fig. 1B) had better peak separation compared with that drawn using Ks_gene_ values (Fig. S1D). Thus, the Ks_block_ frequency distribution was used to identify syntenic blocks derived from the Sol-WGT (Fig. 1B). There were three distinct peaks in the median Ks_block_ distribution (Fig. 1B), consistent with recent tandem/segmental duplications and the two WGT events (Sol-WGT and γ) that the Solanaceae lineage has experienced (Huang et al., 2022; Sato et al., 2012). To identify the gene blocks specifically derived from the Sol-WGT event, we selected two Ks cutoff values corresponding to the intersections between the recent tandem/segmental duplications (exponential distribution) and Sol-WGT (normal distribution on the left), and between the Sol-WGT and the γ event (normal distribution on the right) (Fig. 1B). The first cutoff (t1; Ks = 0.59) excluded 95% of gene blocks arising from recent duplications, while the second cutoff (t2, Ks = 0.91) excluded 90% of gene blocks arising from the γ event, resulting in ∼500 syntenic gene blocks and nearly 5000 syntenic genes, which we designated the Sol-WGT duplicates (Fig. S1B and C).

### Spatial divergence in expression of Sol-WGT duplicates

To understand how evolution shaped gene expression after the Sol-WGT event, we used expression breadth of Sol-WGT duplicates across tissues as a measure of expression divergence across different tissue types. For this analysis, we used publicly available transcriptomes from 23 wild-type tissues, specifically excluding tissues that contain trichomes (hereafter called the baseline tissue set) to minimize confounding effects from trichome-specific metabolite-induced expression biases (Table S1). We focused our analysis on the ∼4,400 Sol-WGT genes (2,200 pairs) that have retained only two copies, allowing for a direct pairwise comparison of extant expression breadth (Fig. 2A). During this analysis we specifically checked if a gene is expressed (TPM ≥ 2) or not (TPM < 2) to determine the gene’s expression breadth across different tissue types (see Materials and Methods). To investigate whether the observed expression divergence pattern deviates from the pattern expected in the absence of selection bias, a simulation was performed (see Materials and Methods). The corresponding Z-score was calculated by comparing the observed extant expression patterns to the randomly expected frequency distribution where a positive Z-score indicates significantly more observed cases than randomly expected, and vice versa (Fig. 2A).

Four main expression divergence patterns stood out. First, for ∼10% of Sol-WGT duplicates, expression across different tissues was biased toward one copy, where one copy exhibited a broader expression breadth (> 18 out of 23 tissues) while the other one displayed a narrower breadth (< 6 tissues) or no expression (Fig. 2B; blue dashed lines). This extreme asymmetrical expression divergence pattern occurred more frequently than expected by chance (Z > 2; Fig. 2C; blue dashed line), indicating that the asymmetrical spatial expression divergence patterns are not due to random drift but likely arise from non-neutral evolution.

The second spatial expression divergence pattern involved paralogous pairs exhibiting nearly identical expression, either across all tissues (394 pairs, ∼18%) or most tissues (one gene in the pair expressed in all the tissues while other gene is expressed in ≥ 90% of tissues; 547 pairs, 25%) (Fig. 2B; black dashed lines). Pairs with nearly identical expression were more common than randomly expected, indicating that expression conservation across tissues is not stochastic, but likely maintained by selection (Fig. 2C; black dashed lines). Functional enrichment analysis indicated that the paralog pairs that retained broad expression are involved in forming multi-unit protein complexes (spliceosome, proteasome, photosystem II; Table S2). Genes encoding these types of proteins, are among the most consistently retained following polyploidization events and show enrichment among dosage-sensitive gene sets (Coate et al., 2011; Edger and Pires, 2009). Thus, it is likely that broadly expressed Sol-WGT paralogs pairs are under dosage-dependent selection. However, this hypothesis needs to be tested by calculating the dosage of these genes and comparing it with the ancestral dosage in further studies.

The third pattern, characterized by symmetrical expression partitioning where both paralogs exhibit similar expression breadth across tissues (either in the same tissues or different tissues), was significantly less prevalent than the random expectation (low Z-score; Fig. 2C; gray dashed line). This pattern captures two distinct evolutionary scenarios: similar breadth within the same tissues, which reflects strict functional conservation, and similar breadth across different tissues, which represents expression subfunctionalization. The scarcity of symmetrical expression partitioning observed in our dataset is likely a result of low statistical and biological probability of both paralogs undergoing parallel modifications in their *cis*-regulatory architecture simultaneously. Because the modification or loss of regulatory modules is inherently stochastic, divergence typically proceeds along an asymmetric trajectory. For instance, the divergence of environmental stress responses among duplicated genes in *Arabidopsis* is highly asymmetric, driven by the differential, non-random loss of specific cis-regulatory elements between paralogs (Zou et al., 2009).

The fourth category consisted of Sol-WGT duplicates that were not expressed in any tissue examined (161 gene pairs, ∼9%; Fig. 2C). To rule out the possibility that these genes may be expressed specifically in the tissue types that were filtered out during the preprocessing of transcriptional data, we re-examined their expression in trichome-containing tissues (n=23), namely leaf tissues, the shoot apical meristem after transition to flowering, green internodes, and stems (Table S1: see tissue category column). We found that 137 out of the 161 gene pairs were also not expressed in any of the 23 additional tissue types (Table S2). This finding suggests that these genes became non-functional after Sol-WGT or that their ancestral gene was not functional.

### Inference of post-Sol-WGT expression gains and losses across tissue types

Expression divergence analysis revealed how the expression patterns of extant Sol-WGT duplicate gene pairs differ from each other. However, it remained to be determined whether these differences are due to differential loss or gain of expression relative to the ancestral gene. To address this, we inferred the ancestral expression state before WGT in each of the 23 baseline tissues/organs using phylogenetically informed ancestral state reconstruction with Bayesian inference methods (Huelsenbeck & Bollback, 2001; see Materials and Methods). Instead of building gene family phylogenies, as is typically done in most ancestral state reconstruction studies, we reconstructed phylogenies of only the Sol-WGT paralogs and their post-WGT tandem duplicates (identified using MCScanX; see Supplementary Methods), rooting each tree with the most closely related paralog that evolved prior to the Sol-WGT event (identified through modeling Ks distributions; see Materials and Methods). This was done for several methodological reasons. First, some gene families (e.g., kinases) predate the evolution of flowering plants making ancestral expression states of those genes in angiosperm-specific tissues (e.g., flowers and fruits) biologically uninterpretable (Munro *et al*., 2022). Second, reconstruction accuracy declines substantially as node age increases due to accumulation of substitutions, duplications, losses, and model violations (Williams *et al*., 2006; Litsios and Salamin, 2012). Third, complex duplication and loss histories in ancient gene families can produce pseudo orthologs, which leads to false ancestral state inference (Litsios and Salamin, 2012). Generating Sol-WGT duplicate-specific phylogenies ensures the robustness of ancestral state reconstructions and that interpretation of ancestral states is done using the Sol-WGT event as the starting point.

Comparison of extant expression patterns with those of the ancestral genes revealed that the Sol-WGT duplicates were significantly more likely to lose expression as posited by Ohno, (1970) (average across tissues: 20%) than to gain it (average: 6%) (*t-*test; P < 0.05). This indicates that the reduction in expression breadth compared with ancestral genes has played a major role in shaping the expression divergence between Sol-WGT duplicates (Fig. S3A and B). Nevertheless, there were also a substantial number of gain events indicating that novel expression patterns could be acquired with a higher than anticipated rate, particularly compared with the rate of sequence substitutions of Sol-WGT duplicates (non-synonymous substitution rate of 0.0009 and synonymous substitution rate of 0.0018) shown in Sato et al. (2012).

Given that the Sol-WGT is known to provide genetic material driving lineage-specific traits, such as fruit color and texture (Sato et al, 2012), we asked whether genes with specific biological functions were more likely to gain or lose expression. To test the hypothesis that expression gains and losses preferentially targeted distinct biological processes rather than occurring uniformly across the genome, we performed a Gene Ontology (GO) enrichment analysis (Fig. S3C and D). Sol-WGT duplicates that gained expression in fruit tissues were enriched with functions associated with sugar metabolism (Fig. S3C and D), such as gluconeogenesis, malate transport, and glucuronosyltransferase activity. Also, Sol-WGT duplicates that gained expression only in fruit tissues were enriched with other GO terms related to fruit traits, such as phosphatidylinositol metabolic process, phosphatidylinositol phosphate kinase activity, alpha−L−arabinofuranosidase activity, and L−arabinose metabolic process (Fig. S3C). Homologs of the genes that gained expression only in fruit tissues also have been shown to regulate the mealy texture of fruits such as apples and pears (Tateishi *et al*., 2005; Nobile *et al*., 2011). Since paralogous genes that gained expression in fruits are enriched in functions related to fruit quality, our findings align with those of Sato et al. (2012), who proposed that the Sol-WGT event contributed to the evolution of favorable fruit traits.

Functional divergence of duplicates is known to be a primary source of trait evolution (Sato et al., 2012). Traits such as fruit quality, inflorescence branching, and flowering time have been key targets during tomato domestication. Therefore, we speculated that domestication-related genes may have gained expression in the tissues related to these traits. To address this, we curated a list of published domestication-associated genes and identified those that originated from the Sol-WGT event (Table S3). We then compared the extant expression patterns of these Sol-WGT duplicates with their reconstructed ancestral expression states to assess whether they gained new expression related to domestication traits after the Sol-WGT event. We found that *Style2.1* (*Solyc02g087850*), which regulates style length and inflorescence branching, and *STEM3* (*Solyc01g092950*), which controls flowering time and inflorescence branching, showed evidence of neofunctionalization, with one paralog showing gain of expression in floral tissues following the Sol-WGT event (Fig. S4, green boxes). Apart from the neofunctionalization events, we also observed subfunctionalization among domestication-associated genes. For example, the *SUN* gene (*Solyc10g079240*), which regulates fruit shape, and its paralog underwent subfunctionalization, with ancestral expression in fruit tissues, stem, root, and callus partitioned between the two duplicates (Fig. S4, blue box). A similar evolutionary fate was observed for another fruit quality-related gene, *Lin5* (*Solyc09g010080*), which regulates fruit sweetness. The ancestral expression pattern was partitioned between fruits, trichome callus, and seedling roots of *Lin5* and its paralogs (Fig. S4, pink box). Other tomato domestication genes, such as *CyscB* (*Solyc04g040190*), *CLAVATA3* (*Solyc11g071810*), *EJ2* (*Solyc03g114840*), and *NOR* (*Solyc10g006880*), have retained the ancestral expression pattern in tissues related to domestication traits (Fig. S4).

### Changes of expression levels during the evolution of Sol-WGT duplicates

Previous studies have shown that changes in gene expression level (i.e., expression dosage of a gene) are deleterious for highly expressed genes (Gout, Kahn and Duret, 2010). Therefore, we wanted to test whether the magnitude of ancestral gene expression affects the evolutionary trajectory of Sol-WGT duplicates. Given our observation that ∼74% of Sol-WGT duplicates have retained the ancestral expression state (expressed or non-expressed) despite 52–91 million years of divergence, we hypothesized that high ancestral expression imposed a selective constraint against dosage changes. To test this, we categorized expression levels into three groups: no expression (TPM < 2, N), low expression (2 ≤ TPM < threshold value, L), and high expression (TPM ≥ threshold value, H), where the threshold value corresponds to one of multiple thresholds that were defined based on the TPM distribution to assess the robustness of our findings (Fig. 3A). We then quantified evolutionary divergence in two complementary ways: first, by calculating the frequency of state transitions (e.g., low ancestral expression state to high extant expression state, L-to-H) for individual paralogs against a null model (Fig. 3B), and second, by measuring the magnitude and direction of expression deviation for both paralogous copies relative to their shared ancestor against a null model (Fig. 3C). To simplify the analysis, we first examined evolutionary divergence in the trichome, the most specialized tissue investigated in this study.

**Figure 3:**
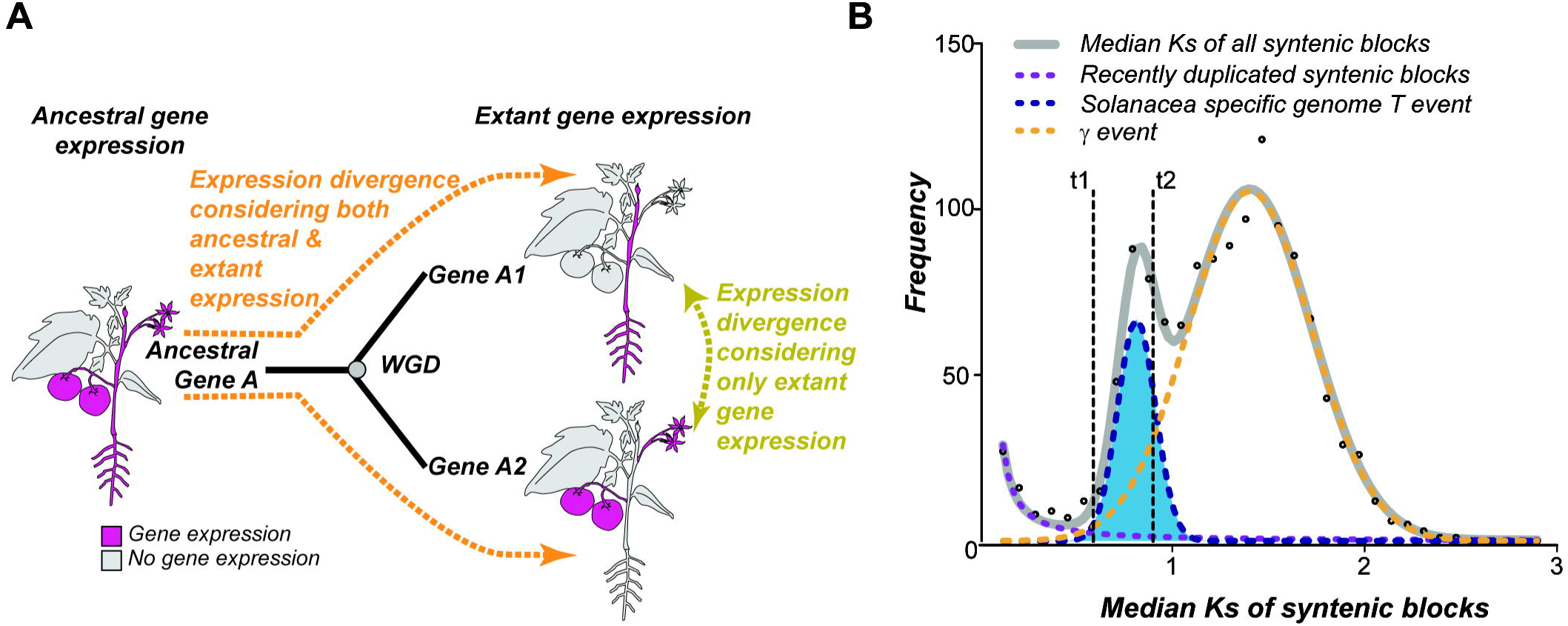
Expression evolution of Sol-WGT duplicates from ancestral expression states in trichomes. **(A)** Modeling of expression level (TPM) distributions of all tomato genes across all tissue types to determine thresholds for categorizing genes as not expressed (N), lowly expressed (L), or highly expressed (H, see Fig. S2). The normal distribution and exponentially decaying distribution separated by mixed modeling are represented by blue and yellow solid lines, respectively. Pink dashed line (at 15^th^ percentile): the threshold for calling a gene “expressed” (TPM ≥ 2); vertical dotted lines: different percentile thresholds for classifying genes as “highly expressed” based on the normally distributed TPM values. The red dashed line (at the 40^th^ percentile) indicates the final TPM threshold used to call a gene highly expressed throughout the study. **(B)** Cartoon illustrating the calculation of Z-scores to assess expression divergence of Sol-WGT duplicates from the ancestral state. The 3×3 table records the observed frequency of each ancestral-to-extant expression state combination. A simulated table was also produced by assigning each gene to any expression state (N, L, and H) independently of ancestral expression state. Z-scores was calculated by comparing observed ancestral-to-extant expression pattern frequencies with the simulated frequency distribution. **(C)** Diagram illustrating six categories of expression deviation. N, L, and H were designated 0, 1, and 2, respectively, and expression deviation between two states was calculated as the difference between these values. For example, the expression deviation between N and H is +2. **(D)** Heatmaps depicting the Z-scores for observed frequencies of expression divergence scenarios for Sol-WGT duplicates calculated comparing observed the changes of the expression levels from ancestral expression to the extant expression and simulated the changes of the expression levels under a null model. Different panels depict the Z score calculation when cutoff for “highly expressed” expression state were implemented. **(E, F and G)** Heatmaps showing Z-scores depicting the expression deviation of Sol-WGT duplicate pairs when the expression state of the ancestral gene was N (E), L (F), or H (G). Z-scores were calculated by comparing the observed frequency of expression deviations between syntenic paralogs to the expected frequency under a random expression model. The expected frequencies were generated through simulations, where extant expression states were randomized. Color indicates Z-score: red indicates higher-than-expected frequencies, and blue indicates lower-than-expected frequencies. **Alt Text:** Multiple panels showing distribution models, a 3×3 state transition table, and heatmaps showing Z-scores for expression level deviations from ancestral states at various threshold percentiles.

In trichomes, we observed higher-than-expected frequencies of conserved expression states, specifically for non-expressed (N-to-N), lowly expressed (L-to-L), and highly expressed (H-to-H) ancestral-extant pairs. This trend of conservation was consistent regardless of the threshold used, particularly when ancestral genes were either non-expressed or highly expressed (*t*-test; *P* ≥ 0.05, Fig. 3D), with the exception of the L-to-L scenario when TPM threshold for highly expressed genes were set to the 25^th^ percentile (TPM = 3.9), as only a small number of genes had expression levels between 2 TPM and 3.9. Approximately 92% of Sol-WGT paralogs retained their non-expressed ancestral states and 72% retained high expression ancestral states in trichomes (Fig. 3D). A similar trend was observed for the deviation of ancestral-to-extant combinations for Sol-WGT duplicate pairs (Fig. 3E). The ancestral expression state was retained when the ancestral gene was either non-expressed or highly expressed. This frequency of this pattern was significantly higher than that expected from a null model, both for the frequency of state transitions for individual paralogs (N-N: ∼76% and H-H: ∼54%; Fig. 3D) and for the expression deviation between paralog pairs (N: ∼76% and H: ∼77%; Fig 3E and G). Conversely, for ancestral genes with low expression (L), only ∼41% of individual paralogs retained the ancestral state, while 36% lost expression and 23% shifted to high expression (Fig. 3D). This trend was further confirmed by analyzing the expression deviation between paralog pairs: only ∼22% of pairs retained low expression in both copies, whereas the majority had asymmetrical (L/N: ∼35%) or loss-gain (N/H: ∼26%) patterns (Fig. 3F). In all three scenarios, these frequencies were higher than null model expectations. This pattern suggests that the low ancestral expression is more likely to be changed during the evolution of Sol-WGT duplicates, indicating weaker selective constraints on lowly expressed genes (Fig. 3F).

Considering that trichomes are specialized tissues, we next evaluated whether the observations about expression divergence hold for root hairs, seeds, and shaved stems, which represent other specialized (root hair) and more general (seed, shaved stem) tissue types (Fig. S5). We observed similar expression evolutionary patterns across these tissue types, indicating that a common mechanism underlies the evolutionary fates of Sol-WGT duplicates in terms of expression. Overall, we observed that Sol-WGT duplicates are more likely to retain their ancestral expression pattern than to lose expression or gain a new expression pattern. Our analysis suggests that transition between expression states is more common when the ancestral gene exhibits low expression.

### Coordinated spatial expression evolution among Sol-WGT duplicates

Our analyses of Sol-WGT duplicates showed that spatial expression divergence is highly asymmetric (Fig. 2B) and that the patterns of expression evolution are similar across different tissue types (Fig. 3, Fig. S5). Under a model where expression in individual tissues evolves independently between paralogs, symmetrical expression divergence would occur only by chance. However, symmetrical expression divergence within tissues was highly prevalent among the analyzed paralogous gene pairs (Fig. 2B). Thus, next we checked whether expression evolves in a coordinated manner between different tissue pairs, which may explain asymmetrical expression divergence. To assess this, we tested the null hypothesis of symmetrical expression partitioning, where each Sol-WGT paralog is expressed exclusively in one of the two tissues, meaning that partitioning of expression between duplicates occurs differently between different tissues. In contrast, the alternative hypothesis posits asymmetric partitioning, where one copy is expressed in both tissues while the other is not expressed in either. Tissue pairs for which there were significantly more gene pairs showing asymmetric expression divergence than symmetric divergence (Fisher’s exact test, see Materials and Methods) were referred to as “correlated tissue pairs” because the expression states of the two tissues in question appear to be co-evolving. In contrast, tissue pairs for which there were significantly more gene pairs showing symmetric expression divergence were referred to as “uncorrelated tissue pairs” (Fig. 4A).

**Figure 4:**
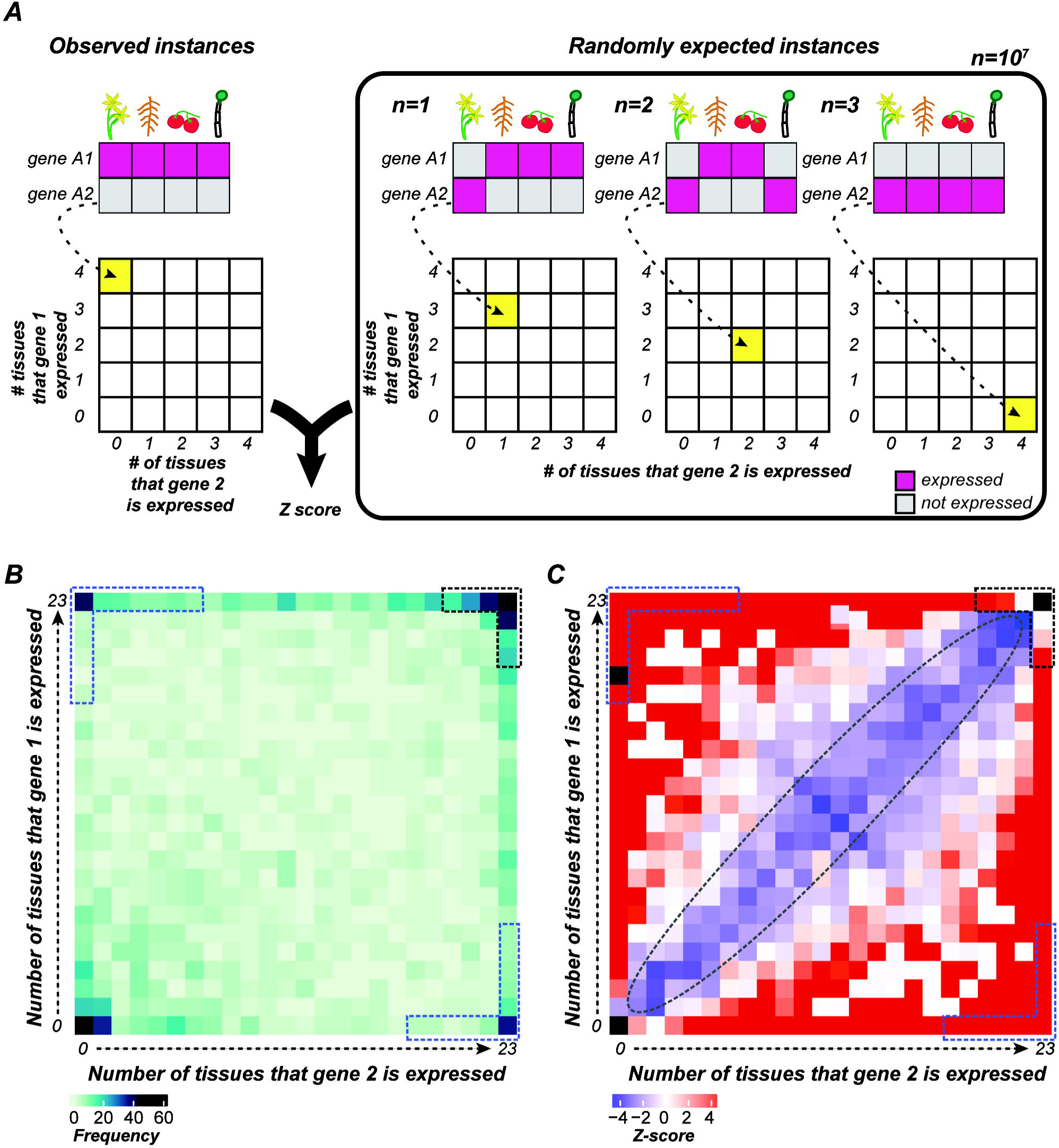
Expression dependency between tissue types for Sol-WGT duplicates. **(A)** Schematic illustrating the calculation of expression association between Sol-WGT duplicates. First, paralogous pairs with at least one broadly expressed gene (expressed in more than 10 tissue types) were excluded to avoid bias. Next, gene pairs showing expression divergence (only one paralogous gene is expressed in a single tissue type) were analyzed for symmetrical or asymmetrical expression divergence. Fisher’s exact test (2 × 2 contingency table is shown in the diagram) was used to test the null hypothesis that there is symmetrical expression divergence between paralogous genes, with log odds ratios calculated as enrichment scores for all possible tissue pair combinations. **(B)** Heatmap depicting the degree and significance of expression coordination between paralogous gene pairs for all possible tissue pair combinations. The upper triangle shows the negative log P-values from Fisher’s exact test; gray boxes represent P-values > 0.05. The lower triangle shows log odds ratios for asymmetrical vs. symmetrical partitioning of paralogous gene pairs, with significant associations marked by dotted block boxes. Dotted vertical and horizontal lines group tissues by similar organ origin. **(C)** Diagram illustrating the evaluation of spatial expression evolution among Sol-WGT duplicates. Paralogous gene pairs were analyzed for coupled gain or loss of expression in each tissue pair. Gene pairs were categorized as “dependently evolving” if one of the paralogs retained the ancestral expression pattern across paired tissues, or “independently evolving” if the ancestral expression pattern was partitioned between two genes in a pair. **(D)** Heatmap showing log odds ratios for dependent vs. independent evolutionary scenarios across tissue pairs. Red denotes tissue pairs with a higher frequency of dependent evolution, while blue indicates tissue pairs with a higher frequency of independent evolution. Tissue pairs with significant associations (q < 0.05) are outlined with dotted boxes. The upper triangle represents expression coordination for paralogs with expression gained in one gene (where the ancestral gene had no expression), and the lower triangle shows associations for paralogs in which ancestral expression was lost in one gene (where the ancestral gene is expressed). Dotted lines group tissues by similar organ origin. **Alt Text:** Conceptual diagrams and heatmaps displaying log odds ratios and significance levels for expression associations and evolutionary gain-or-loss scenarios across tissue pairs grouped by organ origin.

We found that, among all possible tissue pair combinations, the majority tended to be correlated tissue pairs, indicating substantial asymmetrical expression divergence between Sol-WGT duplicates among tissue types (Fig. S6A). This is consistent with the observations for expression breadth divergence, where extreme asymmetrical expression divergence was observed for most Sol-WGT duplicates (Fig. 2B and C). The fact that more than 50% of the paralogous gene pairs had at least one copy expressed in all 23 tissue types analyzed (Fig. 2A) can introduce bias when analyzing tissue pairs. Thus, we asked whether the genes that were preferentially expressed also showed coordinated expression between tissue pairs. We repeated the analysis by filtering out gene pairs with extreme unidirectional expression divergence (pairs with at least one copy expressed in > 10 tissue types). After filtering, significant (q-value < 0.05) coordinated expression was evident among tissues originating from the same organ, particularly among seed tissues, root-derived tissues, and fruit tissues (Fig. 4B).

To further test whether there is coordinated expression evolution between tissues derived from the same organ, even when the genes have a narrow expression breadth, we applied stricter filtering criteria by excluding paralogous gene pairs with at least one copy expressed in > 5 tissue types (Fig. S6B) and paralogous gene pairs for which both genes were expressed in only two tissue types (Fig. S6C). We still observed organ-specific tissue expression correlation among different types of seeds, root-derived tissues, and some fruit tissues, further supporting the hypothesis that organ-specific selective constraints have driven the observed expression divergence of Sol-WGT duplicates. This finding is also consistent with findings in mammals, where post-WGD paralogs have been found to undergo organ-specific expression divergence due to selective constraints and developmental needs (Liao and Zhang, 2006). This expression correlation aligns with the idea that tissues originating from the same anatomical region often exhibit co-regulated gene expression profiles due to shared developmental pathways or regulatory mechanisms (Pattison and Catalá, 2012; Torii, 2021). The co-regulation of gene expression is likely driven by the conservation of cis-regulatory architecture among tissues (Rebeiz, Patel and Hinman, 2015). However, the high frequencies of asymmetrical expression divergence within organs may simply be due to cross-contamination of samples during RNA isolation. This possibility could be ruled out by leveraging data from single-cell RNA-seq studies.

The paralog expression correlation between tissue pairs can occur either by gain or loss of expression in both tissue types. Thus, we tested whether expression correlation between tissue pairs was due to the loss or gain of expression relative to ancestral expression for each post-WGT paralogous gene pair. Gene pairs were classified as “dependently evolving” if one paralog retained the ancestral expression pattern across both tissues while the other diverged. Conversely, genes were classified as “independently evolving” if the ancestral expression pattern was partitioned between the two genes (Fig. 4C). Here, we only analyzed the Sol-WGT gene pairs whose ancestral states were either expressed (but with expression lost in one gene) or not expressed (but expression gained in one gene) for each tissue pair. Fisher’s exact test was used to test the null hypothesis that dependent and independent evolution events were equally distributed across tissue pairs.

Our findings align with the patterns observed for the extant expression coordination where a higher coordinated expression was observed between tissues originating from similar organs (Fig. 4A). Significantly more instances of dependent evolution were identified for gene pairs in which one paralog had either gained or lost expression in both tissues. Notably, the significance of dependent evolution was markedly higher in cases where ancestral expression was lost in one paralog compared to cases of gained expression (Fig. S7E and F). This pattern remained consistent even when the threshold for broadly expressed genes was lowered (Fig. S7A to d; Fig. 4D). For the paralogous gene pairs in which at least one gene was expressed in more than 10 tissue types, there was significant tissue expression dependency for pairs of tissues that originated from the same organ, such as different seed stages, root tissues, and certain fruit tissues. Interestingly, this coordinated expression pattern was only significant for gene pairs in which one paralog had lost the ancestral expression pattern (Fig. 4D). This result suggests that the loss of ancestral expression likely plays a pivotal role in maintaining organ-specific expression patterns. This pattern of reciprocal silencing of gene expression between WGD paralogs has been observed in cotton (Chaudhary *et al*., 2009; Bianconi *et al*., 2018) and in *Arabidopsis* (Liu, Baute and Adams, 2011b). It is likely that selective loss of expression in certain paralogs may preserve critical organ-specific functions necessary for specialized developmental or functional requirements, potentially by maintaining coherence in regulatory networks between tissues with a common origin, facilitating both conservation and diversification of function within organ systems.

## CONCLUSION

In this study, we investigated the evolution of gene expression in tomato following the WGT event, focusing on paralogous gene pairs and the coordination of their expression patterns between tissues. We analyzed our data using two approaches: (1) comparing the divergence of extant expression patterns between Sol-WGT duplicates and (2) analyzing the divergence of ancestral tissue expression patterns between Sol-WGT duplicates.

We demonstrated that asymmetrical divergence of extant expression between paralogous gene pairs is common. Comparing ancestral to extant expression states further revealed that the retention of ancestral expression was more common in Sol-WGT duplicates, particularly when the ancestral gene was either highly expressed or not expressed. Expression gain or loss compared to the ancestral expression state was frequently observed among paralogous pairs whose ancestral genes were lowly expressed, indicating that ancestral expression partitioning might be more prevalent for genes with low ancestral expression levels. We found that genes that gained expression in different tissues were enriched in functions favorable for domestication. From an agricultural perspective, the enrichment of expression gains in fruit-specific pathways illustrates how ancient polyploidy events ‘prime’ plant genomes for subsequent agronomic selection.

We also showed coordinated expression divergence within organ systems in Sol-WGT duplicates. This pattern supports the idea that a common regulatory mechanism governs gene expression in tissues within the same organ type. Organ-specific selective constraints and shared developmental pathways likely play a key role in shaping the expression evolution of paralogous genes, maintaining coherence within organ systems.

By combining comparative analysis of paralogous gene expression and phylogeny-guided ancestral state reconstruction, we were able to generate new insights into the evolutionary dynamics of gene expression following WGD. Our findings highlight the role of selective constraints and tissue-specific pressures in shaping the gene expression landscape of tomato. These findings contribute to a deeper understanding of how WGD influences gene expression patterns and the evolution of plant genomes.

## ACKNOWLEDGEMENT AND FUNDING

We thank Kenia Segura Abá, Brianna Brown, Paulo Izquierdo, Rajneesh Singhal, Jingyao Tang, Edmond Anderson, Thejesh Mallidi, and Shang-Huan Chiang for providing feedback on the figures. This work was funded by the National Science Foundation (IOS 2218206 to T.R., R.E.K., R.L.L., S.-H.S., and M.D.L.-S.) and the National Natural Science Foundation of China (32370241 to PW).

## COMPETING INTERREST

The authors declare that they have no competing interests to influence the work reported in this paper.

## AUTHOR CONTRIBUTIONS

Author Contributions: T.R., P.W., R.L.L., and S.-H.S. designed the research; T.R. and H.C. performed research; T.R. analyzed data; T.R., R.E.K., P.W., M.D.L.-S., and S.-H.S. wrote the paper. All authors have read and approved the final manuscript.

## DATA AVAILABILITY STATEMENT

The custom scripts used in the data processing, analysis and visualization can be found at https://github.com/ShiuLab/Manuscript_Code/tree/master/2025_Tomato_expression_evolution, https://github.com/ShiuLab/RNA-seq_data_processing, and https://github.com/ShiuLab/Ancestral_states_reconstruction.git. All the supplementary methods and figures have been submitted alongside the manuscript.

## Notes

### Competing Interest Statement

The authors have declared no competing interest.

