## Supplemental Materials for "Spatial expression evolution of duplicated genes post-whole genome duplication in cultivated tomato"

**New Phytologist Supporting Information**

**Article acceptance date:**

**The following Supporting Information is available for this article:

Methods S1:
Determination of the cutoff for gene expression**

A TPM > 0 does not always mean that a gene is expressed in a given tissue, as genomic contamination can also map to non-expressed genes and there may be errors in read mapping. Thus, to reduce the number of false positive expressed genes in our dataset, we set a threshold expression level to call whether a gene is expressed by modeling the distribution of TPMs of the genes in 23 tissues using mixed models (<https://github.com/ShiuLab/Manuscript_Code/tree/master/2025_Tomato_expression_evolution>). Based on the modeled distributions, TPM thresholds to call a gene as lowly expressed (2 ≤ TPM < 10) and highly expressed (TPM ≥ 10) were determined (Fig. S1A). The lowly expression threshold of TPM 2 was determined as it eliminated 90% of the false negative expression (S1 Fig B; Yellow distribution) while highly expression threshold which was at 40^th^ percentile of the true positive distribution (S2 Fig blue line) was determined as it eliminated 99% of the false positive expression.

**Identification of paralogous genes derived from the Solanaceae-specific WGT event**

To identify syntenic blocks derived from WGD events, MCScan2 (Wang et al., 2012) was used, with the modified gene annotations from the *Solanum lycopersicum* ITAG 4.0 release as input to define gene locations. Specifically, all-by-all BLASTP analysis was performed for all the tomato genes, with an e-value threshold of 1e-5; a maximum of five subjects per query was kept and fed into MCScan pipeline with default settings, where a syntenic block requires at least five collinear genes between two genomic segments. Using this approach, both syntenic blocks and tandem duplicates were detected, which were used to build phylogenetic gene trees (see section “Phylogenetic tree construction”).

The identified syntenic blocks can be derived from different WGT events or local segmental duplication. Thus, to identify the syntenic gene blocks that originated from the Solanaceae-specific WGT event (Sol-WGT), synonymous substitution rates (Ks) were calculated between syntenic paralogous genes using the MCScan2 pipeline. The median Ks of syntenic gene pairs from each pair of syntenic blocks was also calculated to capture the summarized rate for syntenic blocks. Both the Ks and the median Ks distributions were fit using a mixed model, which contains an exponential decay function and two normal distribution functions (Fig. 1B and Fig. S1D). The normal distributions with median Ks values of 0.71 and 1.35 are indicative of the Solanaceae-specific and γ WGT events, respectively, and syntenic genes within these gene blocks were identified as paralogous genes corresponding to these WGT events.

**Phylogenetic tree reconstruction**

To reconstruct the evolutionary history of specific gene lineages, we generated phylogenetic trees using syntenic Sol-WGT paralogous genes (see section: “Identification of paralogous genes derived from the Solanaceae-specific WGT event”) and any tandemly duplicated genes located immediately adjacent to either syntenic copy (genes immediately upstream and downstream of the Sol-WGT duplicate without any intervening genes). All the genes in a tandem array were included. For example, if Gene A (on Chromosome 1) and Gene B (on Chromosome 5) were identified as a syntenic Sol-WGT pair, and Gene A had a local tandem duplicate (Gene A'), all three genes (Gene A, Gene B, and Gene A') were grouped together for phylogenetic analysis. The outgroup for each phylogeny was selected from syntenic paralogous genes derived from the older γ event.

To determine the optimal outgroup for each syntenic paralog–tandem duplicate group, a pBLAST (Altschul *et al.*, 1990) search was performed between paralogous genes duplicated during the γ event and those from the WGT syntenic paralog–tandem duplicate group. A similarity threshold of 1e-1 was applied, and a maximum of 500 hits per gene were retained. The γ duplicates showing the highest sequence similarity to the WGT syntenic paralog–tandem duplicate genes were selected as potential outgroups. E-values from the pBLAST search were used to measure sequence similarity, and potential outgroups were selected if the E-values between outgroup sequences (γ duplicates) and ingroup sequences (WGT syntenic paralog–tandem duplicates) were higher than those among ingroup sequences. From the potential outgroup candidates, the two sequences with the highest similarity to the ingroup were selected as outgroups for each WGT syntenic paralog–tandem duplicate group.

Subsequently, multiple sequence alignment was performed using protein sequences from both the ingroup and outgroup, utilizing the MAFFT iterative refinement method (Katoh and Standley, 2013). Phylogenetic reconstruction was conducted using RAxML version 8 (Stamatakis, 2006) with 1000 bootstrap replicates and automatic model selection to identify the best evolutionary model for each alignment. After constructing the phylogenetic trees, each tree was evaluated to ensure correct outgroup placement by verifying that the selected outgroups did not cluster with ingroup sequences. If any outgroup was incorrectly placed with the ingroup, the outgroup selection process was repeated using the pBLAST approach described above. This iterative process was continued until all trees had the correct outgroup assignments.

**Ancestral expression state reconstruction**

To investigate how expression of WGT syntenic paralogs evolved in *S. lycopersicum*, we reconstructed ancestral expression states using expression abundance for each gene in specific tissues and the phylogenetic trees generated for the syntenic paralog–tandem duplicate groups (see methods section “Phylogenetic tree reconstruction”) using BayesTraits V3.0.2 (<http://www.evolution.reading.ac.uk/BayesTraitsV3.0.2/BayesTraitsV3.0.2.html>). A Brownian motion model (Pagel, 1999) was applied to account for continuous trait variation over time. Variable rate models were used to account for rate heterogeneity across the trees (Venditti, Meade and Pagel, 2011), and ancestral trait values for internal nodes were estimated using independent contrasts (Felsenstein, 1985; Freckleton, 2012), which is computationally efficient for large datasets (Freckleton, 2012). All models were run in a Bayesian framework, which requires two priors to estimate continuous ancestral traits: the ancestral expression state at the root of the tree (alpha) and the Brownian motion rate parameter (sigma). To determine the best prior ranges for these parameters, a grid search was conducted using 40 example trees with varying ingroup sizes (2, 3, 4, 5; Fig S2). During the grid search, a normal distribution was used for the priors, with the mean values for alpha and sigma ranging from 10 to 200 in increments of 10. Models were built with all combinations of these parameters, and each was run using Markov Chain Monte Carlo (MCMC) chains with a 10% burn-in. A constant seed was implemented to allow comparison across models, and the median marginal log-likelihood of 10 trees of the same size was used to assess model performance, from which the optimal prior range was selected.

To ensure sufficient MCMC chain convergence, we evaluated the Effective Sample Size (ESS) as a measure of parameter convergence. Typically, an ESS > 200 is considered the threshold for adequate chain convergence (Lanfear, Hua and Warren, 2016; Meehan *et al.*, 2018). To validate this threshold, ancestral state reconstructions were performed using a random tree for 10 replicates, running MCMC chains for both 1 million and 10 million iterations. MCMC efficiency was assessed by examining posterior density curves and the ESS for each parameter. Based on this analysis, we selected ESS > 200 as the best criterion for chain convergence in our global ancestral state reconstruction.

Using the optimized model, ancestral expression levels were estimated for all internal nodes in each phylogenetic tree for each of the tissue types. The ancestral state reconstruction for all the paralogous gene families was carried out using custom codes (available at <https://github.com/ShiuLab/Ancestral_states_reconstruction.git>).

**SUPPLEMENTARY FIGURES**

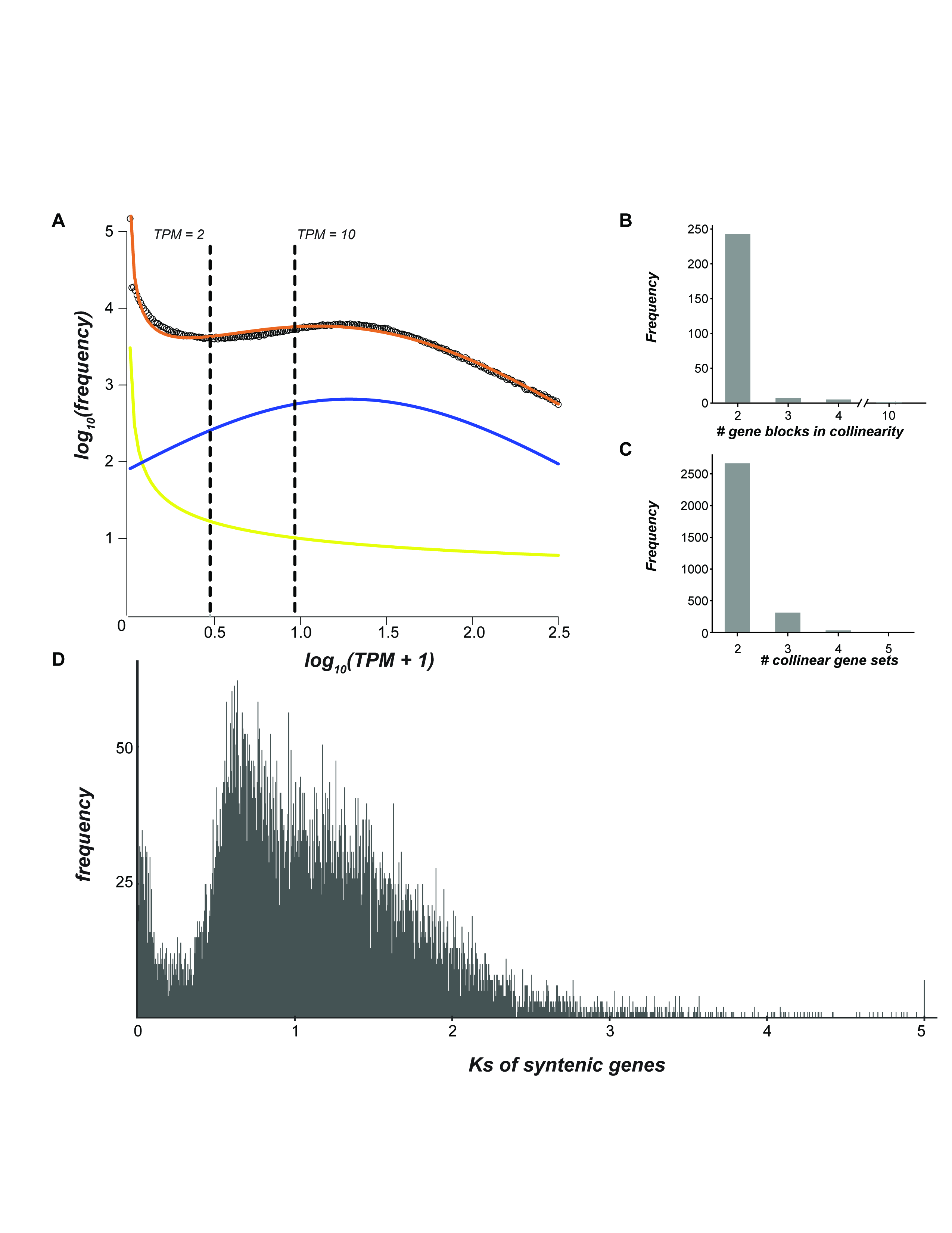

***Figure S1*:** (A) Mixed distribution modeling of gene expression abundance (in TPM). The orange line represents the log_10_ transformed distribution of expression abundance (TPM + 1) for all genes across all the tissue types. The blue line shows the fitted normal distribution, and the yellow line shows the fitted exponentially decaying distribution. The left dotted line indicates the threshold for gene expression, aligning with the 90th percentile of the yellow distribution and the 15th percentile of the blue distribution. The right dotted line represents the threshold for high expression, corresponding to the 99th percentile of the yellow distribution and the 40th percentile of the blue distribution. TPM values for these thresholds are displayed at the top of each dotted line. **(B), (C)** Number of genes and gene blocks sets corresponding to the most recent Solanaceae-specific whole genome triplication event. **(B)** Bar plot showing the frequency of syntenic gene blocks that are in different collinear arrangements that retained after Sol-WGT event. **(C)** Bar plot displaying the frequency distribution of syntenic genes categorized by the number of genes in collinear arrangements **(D)** Distribution of Ks (synonymous substitution rate) calculated between pairs of syntenic paralogous genes.

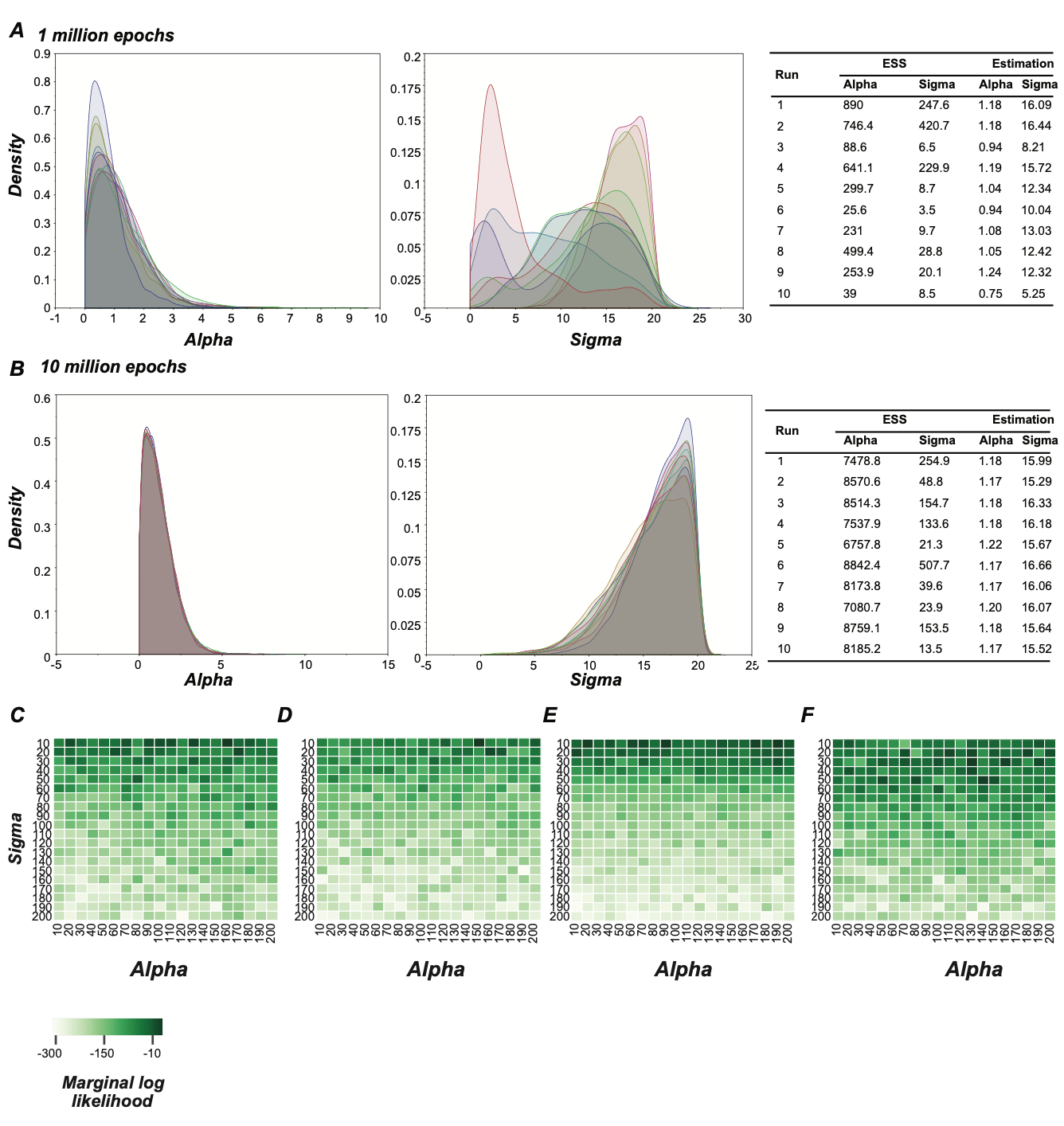

***Figure S2*: Parameter tuning for ancestral state reconstruction. (A), (B)** Plots showing convergence posterior distributions of alpha and sigma parameters for MCMC chains run with 1 million (A) and 10 million epochs of MCMC. The plots on the left and right show the posterior distributions for alpha and sigma parameters, respectively, for 10 different runs. The tables on the right show effective sample size (ESS) after a certain number of MCMC epochs and parameter estimations for alpha and sigma. **(C), (D), (E), (F)** Various prior ranges for sigma (the Brownian motion rate parameter) and alpha (the phylogenetic mean parameter) were tested to build ancestral state models across 10 example trees with ingroup sizes of 2 (C), 3 (D), 4 (E), and 5 (F). Colors represent the median marginal log likelihood across models built on these trees, with darker green indicating higher model performance.

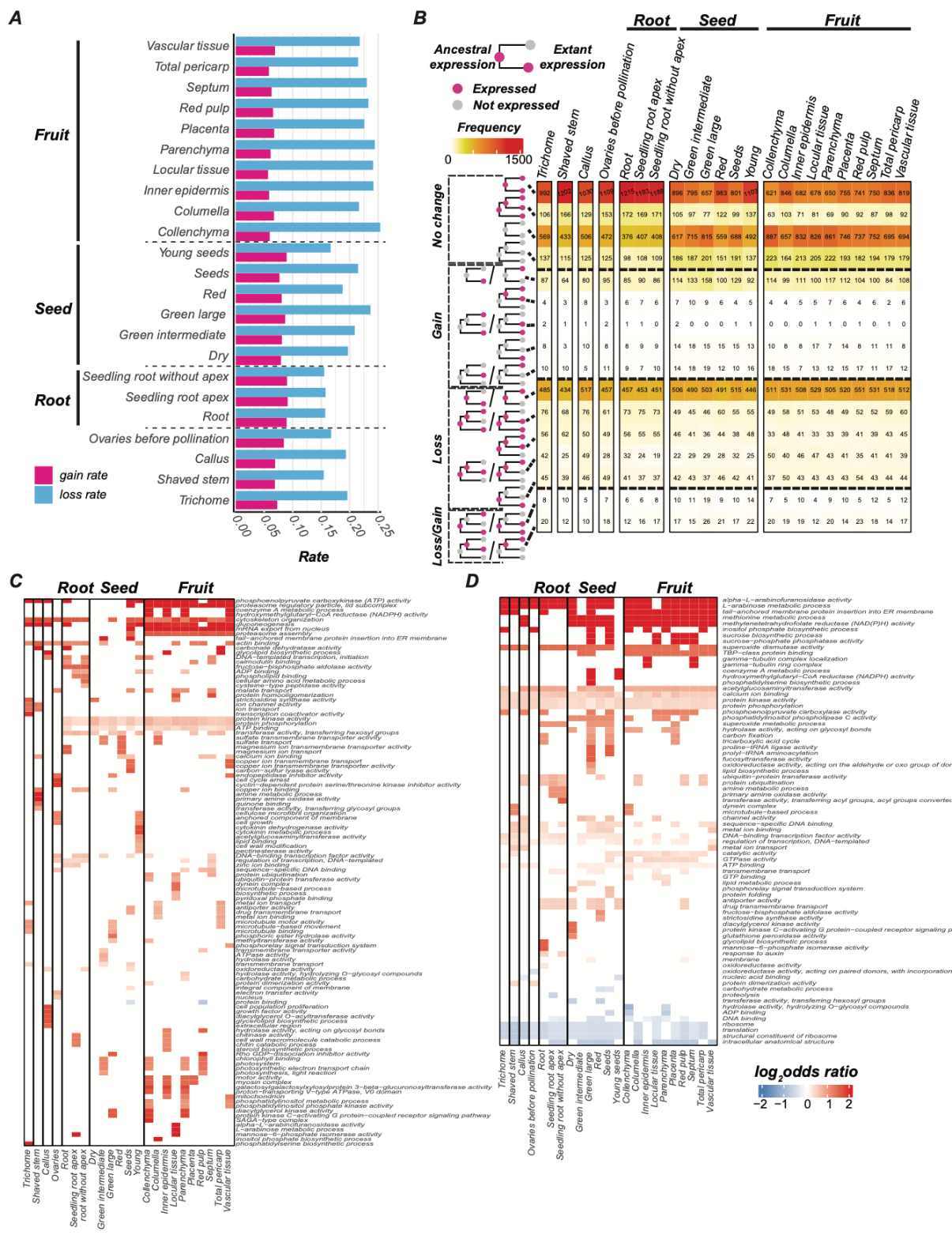

***Figure S3:* Analysis of different expression evolutionary scenarios across tissue types. (A)** Expression gain and loss rates across tissue types following a whole-genome triplication (WGT) event. Red represents expression gain rates, while blue indicates loss rates. The tissue origin is labeled on the left, with similar tissues from the same organ separated by dotted lines. **(B)** Frequency of distinct expression evolution trajectories among syntenic paralogs derived from a WGT event. Rows of the heatmap display combinations of ancestral and extant expression states, categorized by expression changes: gain, loss, both gain and loss, or no change. Colors represent the frequency of each expression evolution type across various tissue types. Columns are grouped by the organ of tissue origin, allowing comparison of evolutionary patterns across different tissues and organs. **(C), (D)** Heatmaps showing log odds ratios for significantly enriched GO terms across the genes that gained (C) and lost (D) ancestral expression in at least one of the syntenic paralogs.

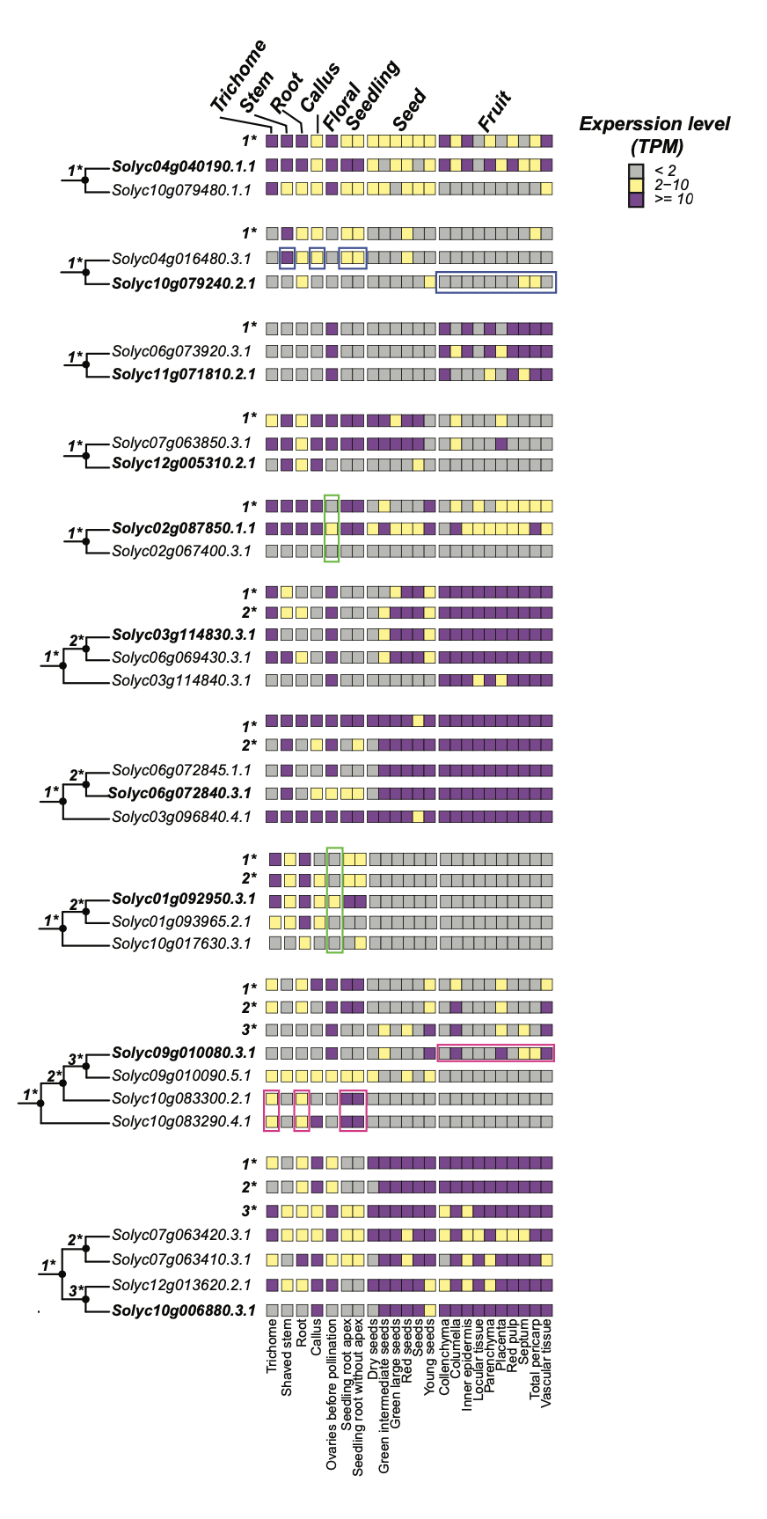

**Figure S4** **Ancestral and extant expression states of tomato domestication genes that were duplicated during the Sol-WGT event.** The phylogeny shows the evolutionary relationships between tomato domestication genes and their paralogs. The heatmap shows categorized extant and ancestral expression levels. *1, *2, and *3 indicate expression levels for ancestral genes at the corresponding nodes of the tree while *1 always corrospod with Sol-WGT event. The green boxes show specific neofunctionalization events and blue and pink boxes highlights different subfunctionalization events.

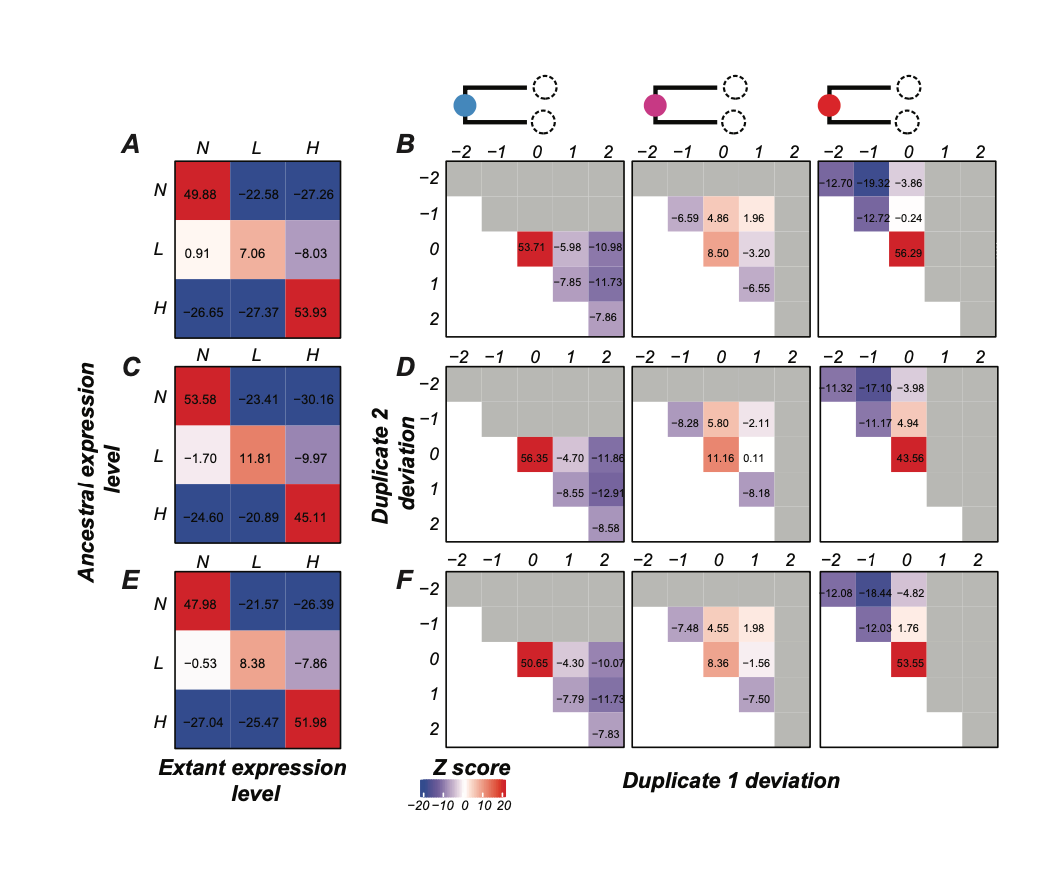

**Figure S5: Expression divergence and deviation of syntenic paralogs derived from the Solanaceae-specific whole genome triplication event.
(A), (C), (E)** Heatmaps illustrate expression divergence patterns for syntenic paralogous genes relative to the ancestral gene in shaved stems (A), young seeds (C), and root hair with apex (E). Z-scores represent observed frequencies compared with random divergence scenarios: red indicates higher-than-expected frequencies, and blue indicates lower-than-expected frequencies. Divergence patterns are shown for different TPM thresholds: H (highly expressed, TPM > 10), L (lowly expressed, TPM ≥ 2 and TPM ≤ 10), and N (non-expressed, TPM < 2). **(B), (D), (F)** Heatmaps showing Z-scores for expression deviations of syntenic paralog pairs based on their ancestral expression states (N, L, or H) in shaved stems. Number in the X and y axis correspond with the number of expression state deviation compared to ancestral gene in the first paralog and the second paralog (B), young seeds (D), and root hair with apex (F). Z-scores were calculated by comparing the observed frequency of expression deviations to the expected frequency under a random expression model.

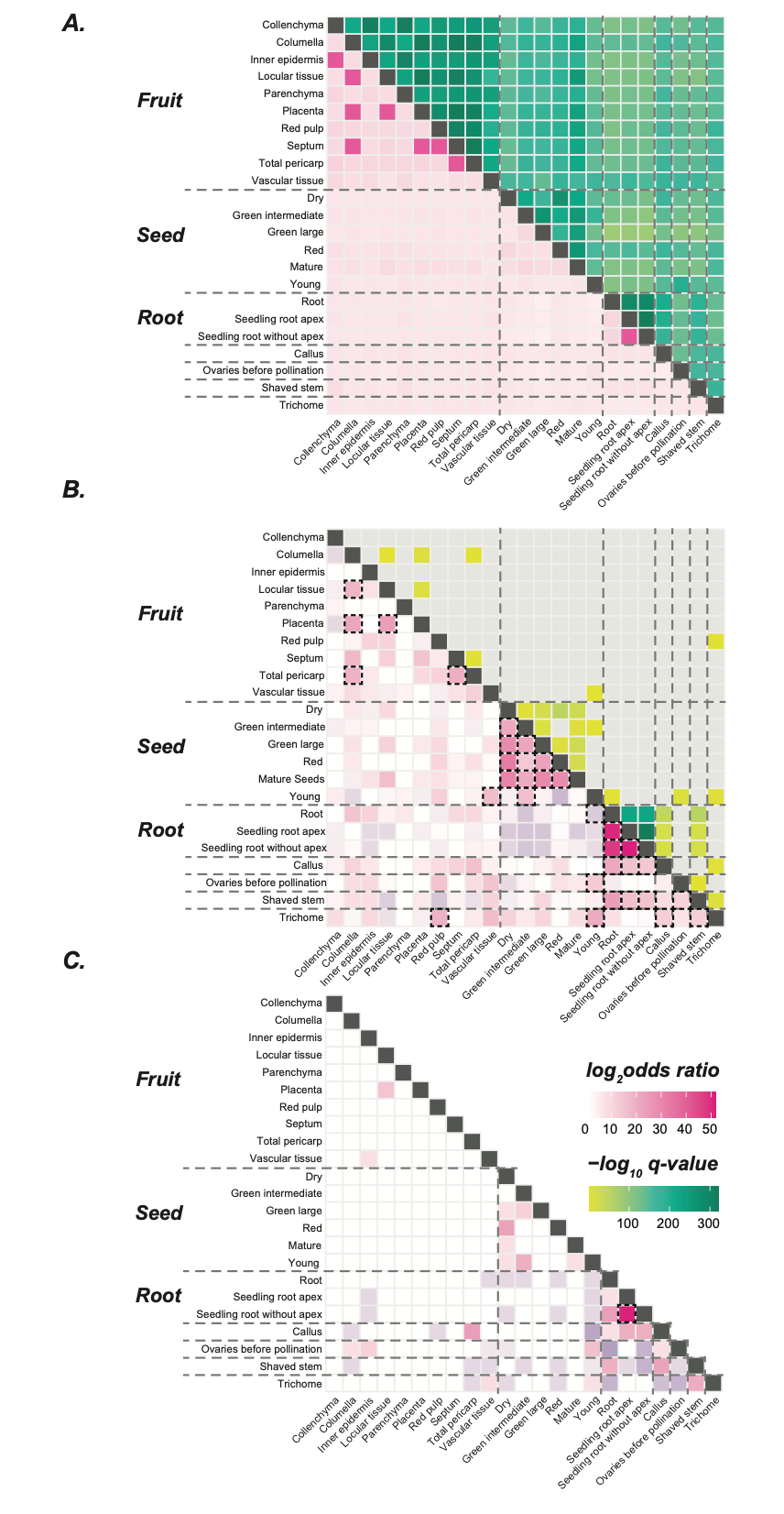

**Figure S6:** **Expression dependency between tissue types for paralogous gene pairs originating from the Solanaceae-specific genome triplication event, analyzed for genes with varying expression breadth. (A)**, **(B)**, and **(C)** Heat maps showing expression associations between pairs of tissues when considering all Sol-WGT paralogous genes that shows an expression divergence at the tissue pair in question(B), paralogous pairs expressed in no more than five tissues (B), and paralogous pairs expressed in no more than two tissues (C). The upper triangle in (C) is omitted, as only one tissue combination showed significant enrichment. The upper triangle displays P-values for expression association between tissue pairs, with colored cells indicating significant asymmetrical expression divergence and gray cells indicating non-significant divergence. The lower triangle shows log odds ratios for asymmetrical versus symmetrical partitioning of paralogous gene pairs, with significant associations marked by dotted boxes. Dotted vertical and horizontal lines group tissues by organ origin.

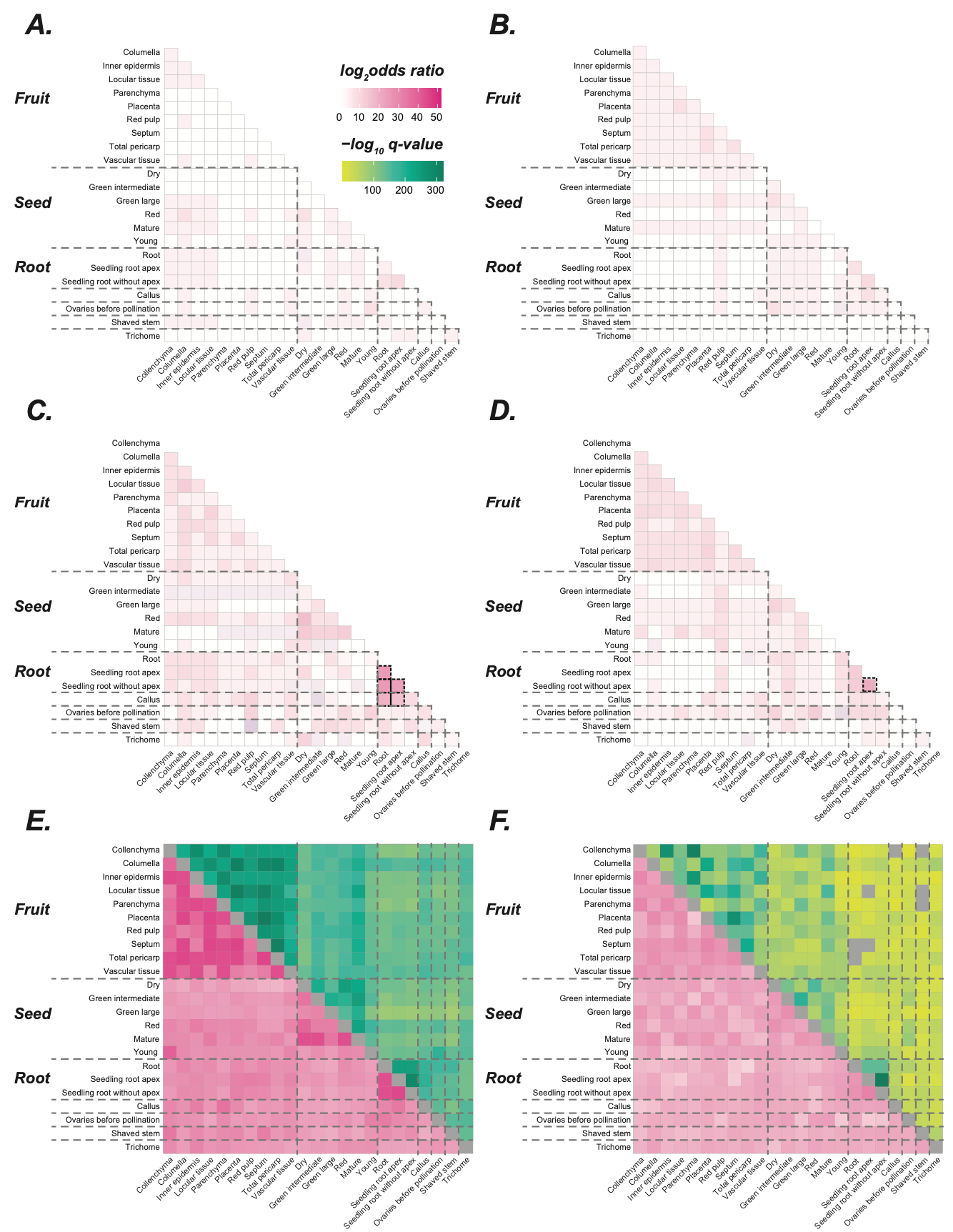

**Figure S7: Coordination of spatial expression evolution in Sol-WGT duplicate genes with varying expression breadth when considering ancestral expression levels. (A)**, **(C)**, and **(E)** Heatmaps representing spatial expression evolutionary associations for paralogous gene pairs that lost ancestral expression in one paralog, across pairs expressed in up to two tissues (A), up to five tissues (C), and all genes (E). **(B)**, **(D)**, and **(F)** Heatmaps showing spatial expression evolutionary associations for paralogous gene pairs with gained expression in one paralog, across pairs expressed in up to two tissues (B), up to five tissues (D), and all genes (F). Heatmap displays log odds ratios for the enrichment of dependent vs. independent evolutionary scenarios across tissue pairs, with red indicating tissue pairs with a higher frequency of dependent evolution and blue showing pairs with more independent evolution. Tissue pairs with significant evolutionary associations across all syntenic paralogs are outlined with dotted boxes, and dotted lines group tissues by organ origin. For cases with over 25% of tissues showing significant evolutionary associations, P-values are shown in the upper triangle, with darker colors indicating stronger significance.

**SUPPLEMENTAL TABLE LEGENDS**

**Table S1:** Metadata for the RNA-seq samples used in this study

**Table S1:** Significantly enriched GO terms from the GO enrichment analysis done for the syntenic paralogous gene pairs that were expressed in all 23 tissues

**Table S3:** Tomato domestication genes that are duplicated from Sol-WGD event
